# Simulation of calcium signaling in fine astrocytic processes: effect of spatial properties on spontaneous activity

**DOI:** 10.1101/567388

**Authors:** Denizot Audrey, Arizono Misa, Nägerl U. Valentin, Soula Hédi, Berry Hugues

## Abstract

Astrocytes, a glial cell type of the central nervous system, have emerged as detectors and regulators of neuronal information processing. Astrocyte excitability resides in transient variations of free cytosolic calcium concentration over a range of temporal and spatial scales, from sub-microdomains to waves propagating throughout the cell. Despite extensive experimental approaches, it is not clear how these signals are transmitted to and integrated within an astrocyte. The localization of the main molecular actors and the geometry of the system, including calcium channels IP3R spatial organization, are deemed essential. However, as most calcium signals occur in astrocytic ramifications that are too fine to be resolved by conventional light microscopy, most of those spatial data are unknown and computational modeling remains the only methodology to study this issue. Here, we propose an IP3R-mediated calcium signaling model for dynamics in such small sub-cellular volumes. To account for the expected stochasticity and low copy numbers, our model is both spatially explicit and particle-based. Extensive simulations show that spontaneous calcium signals arise in the model via the interplay between excitability and stochasticity. The model reproduces the main forms of calcium signals and indicates that their frequency crucially depends on the spatial organization of the IP3R channels. Importantly, we show that two processes expressing exactly the same calcium channels can display different types of calcium signals depending on channels spatial organization. Our model with realistic process volume and calcium concentrations successfully reproduces spontaneous calcium signals that we measured in calcium micro-domains with confocal microscopy. To our knowledge, this model is the first model suited to investigate calcium dynamics in fine astrocytic processes and to propose plausible mechanisms responsible for their variability.

## Introduction

Astrocytes were first characterized as non-excitable cells of the central nervous system since, although they express voltage-gated channels [1], they do not exhibit electrical excitability [2]. Astrocytes excitability instead results from variations of cytosolic calcium concentration [3]. At the cellular level, those calcium signals emerge in astrocytes in response to synaptic activity and may cause the release of molecules called gliotransmitters such as glutamate, ATP, tumor necrosis factor-α, or D-serine, which can modulate synaptic transmission [4–7] and vasoconstriction/vasodilatation [8–11]. This close association of astrocytes to pre- and post-synaptic elements, both structurally and functionally, is referred to as tripartite synapse (see e.g. [12–15] for reviews on tripartite synapses and the associated controversies). On a larger scale, astrocytic calcium signals can modulate neuronal synchronization and firing pattern [16–18] and have been observed *in vivo* in response to external stimuli [19,20]. Altogether, those observations disrupt the traditional view that allocates information processing in the brain to neurons only.

Cell culture, *ex vivo* and *in vivo* studies have demonstrated that astrocytes display both spontaneous calcium signals [19,21–25] and neuronal activity-induced calcium signals [17,20,26]. Astrocytic calcium signals can be localized to synapses [27–30], propagate along processes [31], lead to whole-cell events [32] or even propagate to other cells [33]. Whether this spatio-temporal variability of calcium signals is associated to different physiological functions and whether this could reflect signal integration from different neural circuits is still unknown.

Astrocytic calcium signals are considered to rely mainly on the IP3R calcium channel pathway. Indeed, type-2 IP3R calcium channel is enriched in astrocytes [34] and knocking-out IP3R2 channels abolishes all calcium signals in astrocytic soma and roughly half of them in the cell processes [30]. The molecular origin of the IP3R2-independent signals in processes remains a matter of debate, and could involve calcium fluxes through the plasma membrane [30] and/or other IP3R channel subtypes [35]. At any rate, astrocytes respond to G-protein-coupled receptor (GPCR) agonists with calcium transients [36,37]. The binding of agonists to G_q_/_11_GPCRs activates IP3 synthesis. In turn, the binding of both IP3 and calcium ions to IP3R channels on the membrane of the endoplasmic reticulum (ER) triggers a calcium influx from the ER to the cytosol [38]. The initiation and propagation of calcium signals within astrocytes then relies on the so-called calcium-induced-calcium release (CICR) mechanism: an increase, even small, of the local calcium concentration increases IP3R opening probability thus increasing the probability for local calcium concentration to rise further.

80% of the astrocyte calcium activity *in vivo* take place in the gliapil, which is mostly formed by astrocytic ramifications that cannot be spatially resolved by conventional light microscopy [39], yet account for 75% of the astrocytic volume [40]. According to electron microscopy studies, the perisynaptic astrocyte projections (PAPs) that belong to the gliapil could be as thin as 30-50nm in diameter [41,42]. At this spatial scale, calcium signals are characterized by non-uniform spatial distributions composed of hotspots where calcium signals are more likely to occur and repeat [43,44]. Those observations suggest the existence of subcellular spatial organizations responsible for the spatial distribution of calcium signal patterns. Understanding calcium signaling in PAPs, where astrocytes potentially regulate neuronal information processing, is crucial. However, only calcium signals in thicker processes, around 300nm in diameter, are within reach of current conventional imaging methods [42] and most studies on astrocytic calcium have focused on astrocytic soma and main processes, where characteristics and physiological roles of calcium signals are likely to differ from those of PAPs. Because of the small dimensions and volumes at stake, modeling is currently the only approach that can investigate calcium signal generation, transmission and the effect of spatial properties within PAPs.

Mathematical models of CICR-based signaling date back to the beginning of the 1990s (for recent reviews see e.g. [45–47]). The first IP3R-mediated calcium signaling models assumed perfect mixing of the molecular species and deterministic kinetics (ordinary differential equations) and typically treated IP3 concentration as a parameter [48–50]. In those models, calcium transients emerge as limit-cycle oscillations from a Hopf bifurcation (or a saddle-node on an invariant circle) beyond a critical value of the IP3 concentration. The first astrocyte-specific calcium signaling models arose a decade later. In those models, the IP3 concentration is usually a dynamical variable coupled to calcium but calcium transients still emerge through the Hopf-bifurcation scenario. Notably, those models focused on intercellular IP3 transport within astrocyte networks via gap junctions [51,52]. Stochastic models of IP3R-mediated calcium signaling have also been proposed, that take into account the stochasticity associated with molecular interactions [53–55]. Yet, none of those models take into account the stochastic effects associated with the small volumes and the low copy number of molecules involved in fine processes. Recently, individual-based modeling has been introduced to evaluate the impact of diffusive noise on IP3R opening dynamics [56], but this simplified model disregarded IP3 dynamics and restricted stochasticity to the vicinity of the IP3Rs.

Here, we propose an IP3R-mediated calcium signaling model adapted to the dynamics of CICR in small spatial volumes corresponding to thin PAPs. To account for the stochasticity inherent to small sub-cellular volumes and low copy numbers expected in fine processes, our model is both spatially explicit and particle-based: each molecule is described individually, diffuses in space through a random walk and reacts stochastically upon collision with reaction partners. The kinetics of IP3R channels is accounted for with a simplified version of the 8-state Markov model on which most of the previous CICR models are based. In order to explore the range of dynamical behaviors that the model can display, we first focus on a 2D version of our model, that is less compute-intensive than the 3D one. Extensive simulations of the 2D model show that spontaneous calcium signals arise in the model via the interplay between the excitability of the system and its stochasticity. The model accounts for various forms of calcium signals (“blips” and “puffs”) and their frequency depends on the spatial organization of the IP3R channels. In particular, we demonstrate that the co-localization of sources of calcium influx plays a crucial role in triggering an effect of IP3R clustering on calcium signaling. Finally, as solute concentrations can hardly be defined in 2D, we use a 3D version of the model in order to compare it to experimental data. We show that the spontaneous calcium signals generated by the 3D model with realistic process volume and astrocytic calcium concentrations successfully reproduce the spontaneous calcium transients measured in calcium micro-domains with confocal microscopy in organotypic culture of hippocampal astrocytes. Our model therefore represents the first validated tool to investigate calcium signals in realistic small sub-cellular volumes such as in PAPs, where astrocytes and synapses communicate. This provides a crucial step towards a better understanding of the spatiotemporal response patterns of astrocytes to neuronal activity and beyond, towards astrocyte-neuron communication.

## Materials and methods

### Modeling methods

#### Reaction scheme

The model considers cytosolic calcium and IP3 dynamics in the framework of calcium-induced calcium release (CICR) signaling. The reaction scheme considered is shown in figure 1 A. In short, we consider calcium fluxes between the cytosol and the extracellular space or the endoplasmic reticulum (ER), including via IP3R channels. We also take into account the effect of phospholipase C *δ* (PLC*δ*), that, when activated by calcium, synthesizes IP3. To derive simple models for this scheme, we made the following assumptions:

- We considered that the extracellular and ER calcium concentrations are constant during the simulation, as well as the electrical potentials across the plasma and ER membranes. In this case, calcium outflow from the cytosol to the ER or to the extracellular medium can be lumped into a single first-order rate *α*. Likewise, calcium entry from the extracellular medium or any IP3R-independent Ca^2+^ influx from the ER can be considered constants, too. We lumped them into a single overall constant flux *γ*.
- PLC*δ* enzymes remain located in the cytosol (no translocation) and the amount of their substrate PIP2 is present everywhere in large excess. Under this condition, activated PLC*δ* produces IP3 with constant rate *δ*.

**Fig 1.**
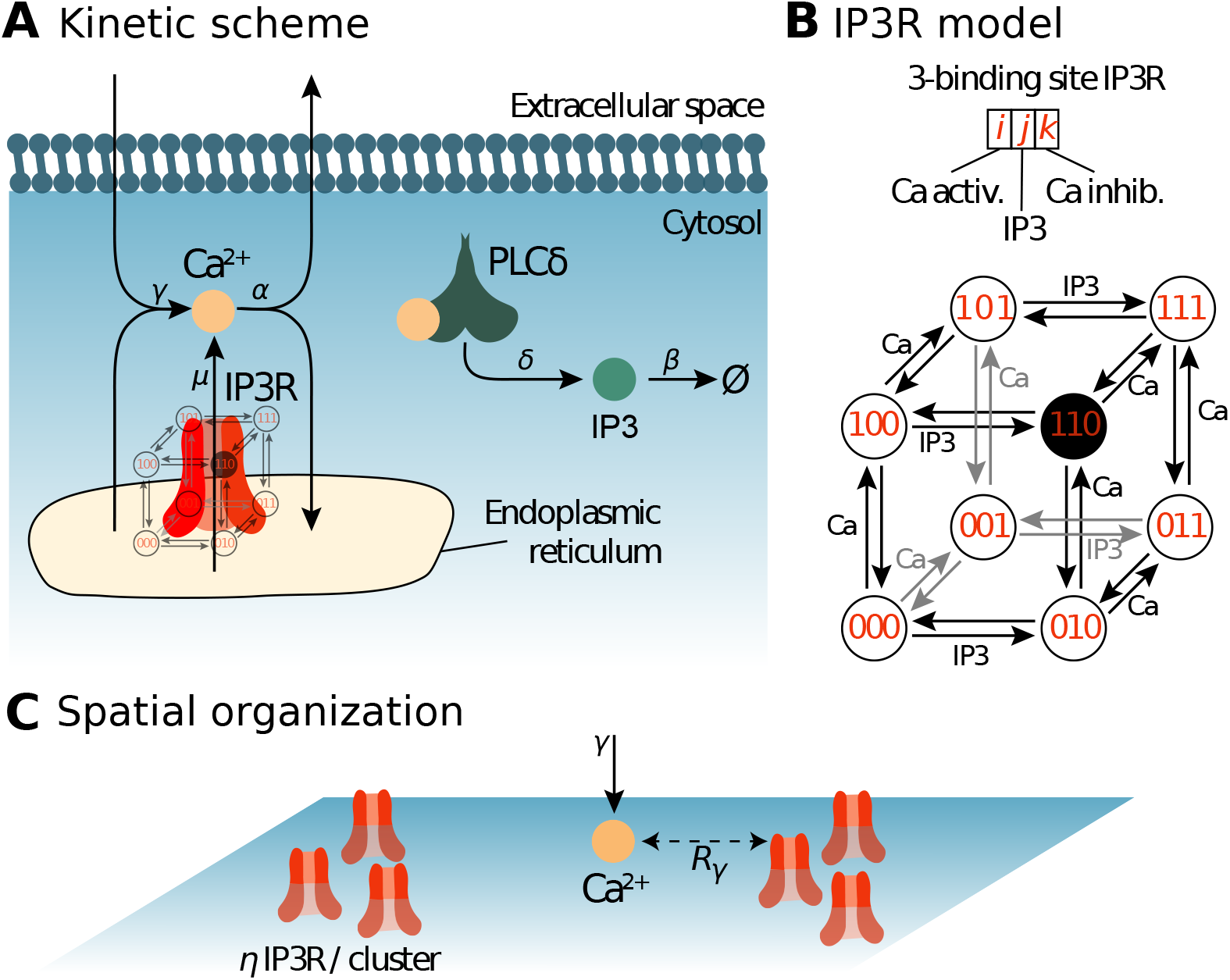
Reaction scheme and IP3R model. The biochemical processes included in the model are illustrated in (*A*): Cytosolic calcium can exit the cytosol to the extracellular space or the endoplasmic reticulum (ER) at a (total) rate *α*, lumping together the effects of ER and plasma membrane pumps. Likewise, Ca^2+^ can enter the cytosol from the extracellular space or from the ER via IP3R-independent flow, with (total) rate *γ*, emulating calcium channels from the plasma membrane. When an IP3R channel opens, calcium enters the cytosol through the channel at rate *μ*. Phospholipase C *δ* (PLC*δ*), once activated by calcium binding, produces IP3 at rate *δ*. Like Ca^2+^, IP3 can bind the IP3R channel and is removed with rate *β*. (*B*) Our model of the kinetics of the IP3R channel is an 8-state Markov model adapted from [48,57]. Each IP3R channel monomer is associated with 3 binding sites, two calcium binding sites and one IP3 binding site. Occupancy states are designated by a triplet {*i,j, k*} where *i* stands for the occupation of the first Ca binding site (*i* = 1 if bound, 0 else), *j* for that of the IP3 binding site and *k* for the second Ca site. The first calcium binding site has higher affinity than the second. The open state is state {110}, where the first Ca and the IP3 sites are bound but not the second Ca site. (*C*) Spatial parameters for the particle-based model. The *N*_IP3R_ IP3R molecules are positioned within uniformly distributed clusters, with *η* IP3R in each cluster. Hence *η* = 1 corresponds to uniformly distributed IP3R (no clustering), while the degree of clustering increases with *η* (for constant total IP3R number). To account for potential co-localization between IP3R-dependent and IP3R-independent calcium sources, the influx of IP3R-independent calcium (at rate *γ*) occurs within distance *R_γ_* of an IP3R. Thus, low values of *R_γ_* emulate co-localization between IP3R-dependent and IP3R-independent Ca^2+^ influx sources.

IP3R channels are gated both by calcium and IP3, with a bell-shaped dependence of the open probability to calcium concentration [57]. To model their dynamics, we used the classical 8-state Markov model proposed in [48,57], with two calcium binding sites and one IP3 binding site for each IP3R (see figure 1B). However we used the following simplifications:

- We considered that the binding or unbinding rate constant of a given binding site is independent from the occupancy state of the other sites (no intra-channel cooperativity). Under this assumption, the rate constant for calcium binding at the first calcium binding site, *a*_1_, does not depend on whether the other two binding sites are bound or not. Thus, the rate constant for {000} + Ca → {100} has the same value as e.g. the reaction {011} + Ca → {111} (where the triplet notation corresponds to the one defined in figure 1). Likewise, the rate constants for Ca^2+^ or IP3 binding or unbinding to the three sites were considered independent from the other occupancy states.
- The open state is assumed to be state {110} (first Ca site and IP3 bound, second Ca site free), as in [48,57]. These latter models further assume inter-channel cooperativity, where IP3R channels assemble as tetramers of which at least three monomers must be in the open state for calcium to be transferred. Here we neglected inter-channel cooperativity and considered that every single channel was open when in the open state i.e., as long as an IP3R channel is open, it is assumed to inject calcium in the cytosol at constant rate *μ*.

#### Monte Carlo simulations of the spatially-explicit stochastic particle-based model

We first modeled the kinetic scheme described in figure 1 with a lattice-free spatially-explicit stochastic particle-based model, referred to as “Particle-based” model below, in two spatial dimensions, with reflective boundary conditions. Each molecule of the system was explicitly modeled with its associated position in space. PLC*δ* and IP3R molecules were considered immobile whereas Ca^2+^ and IP3 molecules were mobile by diffusion. At the beginning of each Monte-Carlo (MC) simulation of this model, the space coordinates for each Ca^2+^, IP3 and PLC*δ* molecules are chosen uniformly at random.

To determine the positions of the *N*_IP3R_ IP3R molecules, we first chose the centers of *N*_c_ = *N*_IP3R_/*η* IP3R clusters uniformly at random in the reaction space, where *η* is the number of IP3R per cluster (as illustrated in figure 1 C). For each cluster, we positioned *η* IP3R molecules uniformly at random within a distance *R_c_* of the cluster center, with 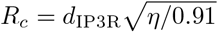, where *d*_IP3R_ is the interaction distance of the IP3R, i.e. the maximal distance between IP3R center and a Ca^2+^ or IP3 molecule below which binding can occur. According to this algorithm, *η* =1 corresponds to randomly distributed independent IP3R molecules (no clustering) whereas IP3R molecules become increasingly clustered when *η* increases, with constant IP3R density within the clusters and constant total IP3R number in the reaction space.

Each MC stimulation step (of duration Δ*t*) consists in iterating the following steps:

1. *Diffusion.* The position of each mobile molecule (Ca^2+^ and IP3) is updated independently according to Brownian motion: 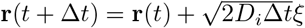, where *D_i_, i* = {Ca, IP3} is molecule *i* diffusion coefficient and *ξ* is a vector of i.i.d. Gaussian-distributed random numbers with zero mean and unit variance. In a subset of simulations, the new position of each mobile molecule was chosen at random in the reaction volume, i.e. **r**(*t* + Δ*t*) = *ζ* were *ζ* is a vector of i.i.d. random numbers uniformly distributed in [0, *L*], with *L* the length of the spatial domain. We refer to this setting as “infinite” diffusion coefficients, *D* = ∞.
2. *Binding.* For each Ca^2+^ ion close enough to a PLC*δ* to react (i.e. when the distance between both is less than the interaction radius of PLC*δ*), a new IP3 molecule is created with probability *δ*Δ*t* at the position of the PLC*δ* molecule. Likewise, each Ca^2+^ or IP3 molecule close enough to an IP3R molecule (i.e. within its interaction radius) can bind it depending on its occupancy state. If the IP3 binding site is free, an IP3 molecule binds with probability *a*_2_Δ*t*. If one of the Ca sites is free, a Ca^2+^ ion binds the free site with probability *α*_1_Δ*t* (first Ca site) or *a*_3_Δ*t* (second Ca site). If both Ca sites are free, binding occurs to the first site with probability *a*_1_Δ*t* and to the second one with probability (1 – *α*_1_Δ*t*)*α*_3_Δ*t*.
3. *Unbinding.* Each IP3R molecule releases its bound Ca^2+^ or IP3 molecules independently, with probability *b*_1_Δ*t* (first Ca site), *b*_2_Δ*t* (IP3 site) and *b*_3_Δ*t* (second Ca site). Ca^2+^ or IP3 molecules that bound the IP3R at the previous (binding) step of the current time step do not unbind.
4. *Removal.* Free cytosolic Ca^2+^ and IP3 molecules are removed from the cytosol with probability *α*Δ*t* and *β*Δ*t*, respectively. Ca^2+^ and IP3 molecules that unbound from IP3R at the previous (unbinding) step of the current time step are not removed.
5. Ca^2^ + *Influx.* For each IP3R channel in the open state {110}, a new Ca^2+^ ion is created in the cytosol at the IP3R position with probability *μ*Δ*t*. A new calcium ion can also be created in the cytosol with probability *γ*Δ*t*, mimicking Ca^2+^ influx from IP3R-independent sources in the ER membrane or through the plasma membrane. Note that the position of this new calcium is not uniform over space but depends on parameter *R_γ_* and works as follows: an IP3R molecule is chosen (uniformly) at random and the new Ca^2+^ ion is positioned uniformly at random within distance *R_γ_* of the chosen IP3R. Therefore low values of *R_γ_* emulate co-localization between IP3R-dependent and IP3R-independent Ca^2+^ influx sources, whereas the location of IP3R-independent Ca^2+^ influx is uniform over the reaction volume when *R_γ_* becomes as large as the volume side length.

Table 1 gives the parameter values used in our 2D simulations, including the initial numbers of Ca^2+^, PLC*δ*, IP3 and IP3R molecules.

**Table 1.** Parameter values and initial conditions of the 2D model. a.u: arbitrary unit. In 2d, by definition, a MC time unit is 100 Δ*t* and one MC space unit is set by the interaction radius of IP3R, i.e. *d*_IP3R_ = 1.0 MC space unit. *δ, β, μ, γ*, *b*_1_, *b*_2_ and *b*_3_ are first order constants, in (MC time unit)^-1^. Diffusion coefficients *D*_Ca_ and *D*_IP3_ are expressed in (MC space unit)^2^.(MC time unit)^-1^ whereas *α, a*_1_, *a*_2_, *a*_3_ are expressed in (MC space unit)^2^.(MC time unit)^-1^.

| Parameter | Description | Value in 2d model |
| --- | --- | --- |
| V | Cell volume | $200 \times 200$ a.u. |
| IP3 dynamics |  |  |
| $IP_0$ | Initial IP3 number/conc. | 15 molec. |
| $D_{IP3}$ | IP3 diffusion | 10 a.u |
| $N_{plc}$ | PLCδ number/conc. | 1000 molec. |
| $\delta$ | PLCδ max rate | 0.1 a.u |
| $\beta$ | IP3 decay | 0.01 a.u |
| Ca<sup>2+</sup> dynamics |  |  |
| $Ca_0$ | Initial Ca<sup>2+</sup> number/conc. | 50 molec. |
| $D_{Ca}$ | Ca<sup>2+</sup> diffusion | varied |
| $\mu$ | Ca<sup>2+</sup> flux through open IP3R | 50 a.u |
| $\gamma$ | cytosolic Ca<sup>2+</sup> influx | 50 a.u |
| $\alpha$ | Ca<sup>2+</sup> decay rate | 1.0 a.u |
| IP3R |  |  |
| $N_{IP3R}$ | IP3R number | 1000 molec. |
| $d_{IP3R}$ | IP3R interact. distance | 1 space unit |
| IP3R binding |  |  |
| $a_1$ | First Ca | 1.0 a.u |
| $a_2$ | IP3 | 1.0 a.u |
| $a_3$ | Second Ca | 0.1 a.u |
| IP3R dissociation |  |  |
| $b_1$ | First Ca | 0.1 a.u |
| $b_2$ | IP3 | 0.1 a.u |
| $b_3$ | Second Ca | 0.1 a.u |

#### Mean-field (MF) dynamics of the perfectly stirred model

With infinite diffusion, the dynamics of the system can be assumed to be perfectly stirred. With that mean-field (MF) assumption, the temporal dynamics of reaction scheme figure 1 can be modeled using ordinary differential equations based on the mass-action law. IP3R dynamics in these conditions can be described with seven ODEs that express the temporal dynamics of the concentration of IP3R in state {*ijk*}, [*ijk*]:

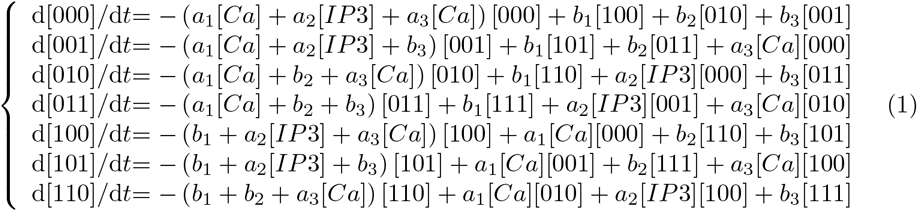

where [*Ca*] and [*IP*3] denote Ca^2+^ and IP3 concentration, respectively. The concentration of the eighth occupancy state, {111} is obtained from conservation of the IP3R, i.e.[1111] = *N*_IP3R_/V – ([000] + [001] + [010] + [011] + [100] + [101] + [110]). IP3 dynamics in the mean-field model is given by:

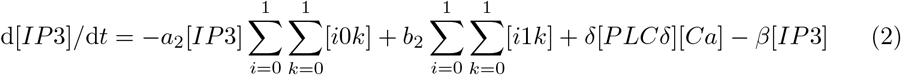

where [*PLCδ*] = *N*_plc_/*V*. Finally, the mean-field dynamics of the free Ca^2+^ is obtained with:

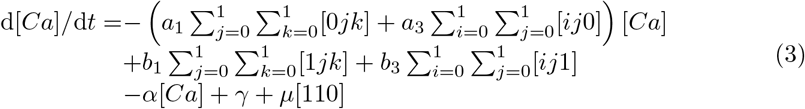

For comparison with the output of the other models, the concentrations were transformed into numbers of molecules by multiplication by the reaction volume *V*.

#### Perfectly-stirred stochastic temporal dynamics (SSA)

For comparison, we also modeled the reaction scheme depicted in Figure 1 using Gillespie’s exact Stochastic Simulation Algorithm (SSA) that accounts for stochasticity due to low copy numbers and assumes perfect mixing of the reactants [58, 59]. Here, the dynamic variables are the number of Ca^2+^ and IP3 molecules in the system, *N*_Ca_ and *N*_IP3_ and the number of IP3R channels in state {*ijk*}, *N*_ijk_. The rates of all the reactions of the scheme of figure 1 are then calculated according to mass-action laws like in the MF model of eq. (1,2,3). For instance, at reaction time t, the rate of reaction {001} + *Ca* → {101} is given by *a*_2_/*VN*_001_(*t*)*N*_Ca_(*t*). The next reaction time *τ* is sampled from an exponential distribution with mean 1/*R_T_*, where *R_T_* is the sum of the reaction rates of all reactions. The next reaction to occur at time *t* + *τ* is chosen as an integer random variable with point probability given by the ratio of its rate to *R_T_*. For instance, for the reaction illustrated above, the probability that this reaction is the one occurring at time *t* + *τ* is *a*_2_/*VN*_001_(*t*)*N*_Ca_(*t*)/*R_T_*. Finally, the variables are updated according to the chosen reaction. In the data presented below, we have modeled each receptor individually, i.e. for each receptor 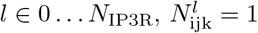 if receptor *l* is in state *ijk*, 0 else. If the illustration reaction described above on receptor *l* is chosen, this means 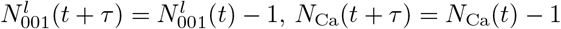 and 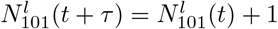. The other variables keep their values.

#### Realistic simulations in 3d astrocytic processes

In order to simulate calcium dynamics within a more refined 3d geometry with realistic volumes and concentrations, we built a model in STEPS (http://steps.sourceforge.net/). STEPS is a software for voxel-based stochastic reaction-diffusion simulations within complex 3d geometries that simulates stochastic chemical reaction-diffusion with a spatialized version of Gillespie’s SSA, usually referred to as the reaction-diffusion master equation (RDME). In RDME, space is partitioned into voxels inside which perfect mixing is assumed, while diffusion between adjacent voxels is modeled as first order reactions [60,61]. STEPS uses a derivative of the SSA in tetrahedral voxels that allows for a better resolution than the cubic voxels mostly used in voxel-based models [62].

##### Geometry

The main advantage of STEPS in the context of the present study is its automatic handling of external and internal membranes [63]. Moreover, STEPS simulations can easily be parallelized [64], a crucial property given the computational burden of such compartmentalized 3d simulations. This allowed us to explicitly describe the presence of the ER membrane inside the 3d cell cytoplasm and the fact that IP3R channels are located in the ER membrane (see figure 6B). The geometry of the reaction volume consisted in a cylinder of length *L*_astro_=1 *μ*m and radius *R*_astro_=0.1 *μ*m. The ER was modeled as a second cylinder, internal, with length *L*_ER_=0.75 μm and radius *R*_ER_=0.03 *μ*m. The resulting cytosolic volume (2.81 × 10^-17^ L) was meshed with 11345 tetrahedra of individual volume 2.48 × 10^-21^*L*, thus ensuring the well-mixed subvolume condition [62].

##### Reactions

In this spatial configuration, we modeled the IP3R-mediated calcium signaling kinetic scheme of figure 1 with IP3R channels positioned on the intracellular ER membrane and according to two model variants:

1. A first variant, referred to as the “No-GCaMP” model, did not include fluorescent calcium indicators. In this 3d model, parameter values were taken, whenever possible, from the literature (Supplementary Table S1). *γ* and *α* values were adjusted to yield basal calcium concentration around 120 nM [65,66]. Likewise, *β* and *μ* were adjusted for a basal IP3 concentration of 120nM [67]. Note that this value is based on recent, precise measurements of IP3 concentration and differs by an order of magnitude from IP3 concentration values routinely used in IP3R-mediated calcium models [68–70]. IP3R density on the ER surface has been measured from TIRF-microscopy analysis in cell cultures [71], reporting IP3R cluster diameters of 0.3 *μ*m at most, with up to 10 IP3R per cluster. The ER surface area in our model is 0.69 *μ*m^2^. Ignoring the potential unclustered IP3Rs [72], this represents a maximum of 4 clusters, thus at most 40 IP3R. We thus set the number of IP3R in our model to 50 channels on the ER surface. Finally, Ca^2+^ and IP3 binding and dissociation constants to IP3R were adjusted to fit our experimental data of calcium micro-domains in organotypic cultures of hippocampal astrocytes.
2. A second variant of the 3d model, referred to as the “GCaMP” model, was obtained by adding GCaMP6s calcium indicators in the cytosol. GCaMP6s are ultrasensitive calcium indicators that fluoresce when bound to Ca^2+^. The fluorescence signal from experimental data indeed corresponds to the concentration of calcium-bound GCaMP6s, which can be quite different from free cytosolic Ca^2+^ trace. Here again Ca^2+^ and IP3 binding and dissociation constants to IP3R were adjusted to fit our experimental data. The parameters related to GCaMP were taken from the available experimental literature and shown in Table 2.

**Table 2.** Parameter values and initial conditions of the 3d GCaMP model. The parameter values for the 3d model listed here correspond to the GCaMP model. The parameter values for the 3d model devoid of GCaMP are presented in Supplementary Table S1. Parameter values in the 3d model have been adjusted to optimize the match with experimental data as described in the Methods section and shown in Fig 6. Note that the values for calcium and IP3 binding or unbinding to IP3R, i.e. the *a_i_*’s and *b_j_*’s parameters below, are smaller in our model than in the literature, probably because our model is not cooperative. For GCaMP6s, we used the diffusion coefficient of calmodulin. The initial number of Ca^2+^ ions was adjusted so that the measured basal GCaMP6s-Ca concentration was around 300nM [65,66].

| Parameter | Description | Value in 3d GCaMP model | Reference |
| --- | --- | --- | --- |
| V | Cell volume | $2.81 \times 10^{-17}$ L | [73] |
| IP3 dynamics |  |  |  |
| $IP_0$ | Initial IP3 number/conc | 3 molec. i.e 177 nM | [67] |
| $D_{IP3}$ | IP3 diffusion | $223 \mu\text{m}^2.\text{s}^{-1}$ | [74] |
| $N_{\text{plc}}$ | PLCδ number/conc. | 1696 molec. i.e 100 $\mu\text{M}$ | [75] |
| $\delta$ | PLCδ max rate | $1 \text{ s}^{-1}$ | - |
| $\beta$ | IP3 decay | $1.2 \times 10^{-4} \text{ s}^{-1}$ | - |
| $\text{Ca}^{2+}$ dynamics | | | |
| $Ca_0$ | Initial $\text{Ca}^{2+}$ number/conc. | 5 molec. i.e 295 nM | [65] |
| $D_{\text{Ca}}$ | $\text{Ca}^{2+}$ diffusion | $13 \mu\text{m}^2.\text{s}^{-1}$ | [74] |
| $\mu$ | $\text{Ca}^{2+}$ flux through open IP3R | $5 \times 10^3 \text{ s}^{-1}$ | - |
| $\gamma$ | cytosolic $\text{Ca}^{2+}$ influx | $8.5 \times 10^{-7} \text{ s}^{-1}$ | - |
| $\alpha$ | $\text{Ca}^{2+}$ decay rate | $5 \text{ s}^{-1}$ | - |
| GCaMP6s |  |  |  |
| $C_{\text{GCa}}$ | GCaMP6s conc. | 169 molec. i.e 10 $\mu\text{M}$ | [76,77] |
| $D_{\text{GCaMP}}$ | GCaMP6s diffusion | $50 \mu\text{m}^2.\text{s}^{-1}$ | [78] |
| $Gcamp_f$ | GCaMP6s Ca binding rate | $2.7 \times 10^5 \text{ M}^{-1}.\text{s}^{-1}$ | [79] |
| $Gcamp_b$ | GCaMP6s-Ca dissociation rate | $1.12 \text{ s}^{-1}$ | [79] |
| IP3R |  |  |  |
| $N_{\text{IP3R}}$ | IP3R number | 50 molec. | [71] |
| $d_{\text{IP3R}}$ | IP3R interact. distance | 1 mesh triangle | - |
| IP3R binding |  |  |  |
| $a_1$ | First Ca | $1.5 \times 10^6 \text{ M}^{-1}.\text{s}^{-1}$ | - |
| $a_2$ | IP3 | $1.5 \times 10^6 \text{ M}^{-1}.\text{s}^{-1}$ | - |
| $a_3$ | Second Ca | $1 \times 10^5 \text{ M}^{-1}.\text{s}^{-1}$ | - |
| IP3R dissociation |  |  |  |
| $b_1$ | First Ca | $70 \text{ s}^{-1}$ | - |
| $b_2$ | IP3 | $70 \text{ s}^{-1}$ | - |
| $b_3$ | Second Ca | $70 \text{ s}^{-1}$ | - |

#### Simulations code

The code of our ODE, Gillespie, Particle-based and STEPS models is available on ModelDB at https://senselab.med.yale.edu/modeldb/enterCode.cshtml?model=247694. The code will be in public access once the paper will be published.

#### Peak detection and analysis

Automated peak detection from the model simulations was based on the statistics of baseline calcium trace. A histogram of Ca^2+^ trace was built with a bin size of 0.25 ions and the mode of this histogram was used to define baseline calcium. A peak initiation corresponded to the time step where calcium trace overcame a peak threshold defined as baseline + *nσC_a_* where *σC_a_* is the standard deviation of the above histogram. The value of n varied between 2 and 4 and was set by hand for each simulation, depending on its signal/noise ratio. The peak was considered terminated when the calcium trace decreased again below peak threshold. This implies that in case of a second calcium peak starting before the first one terminated, both events were considered as being part of the same peak. Peak duration was defined as the time between peak initiation and termination. Peak amplitude was defined as the maximum number of calcium ions reached during the peak duration. In the 3D model, the peak amplitude A was rescaled to facilitate comparison with experimental data, using Δ*F/F* = (*A* – *Ca*_baseline_)/*Ca*_baseline_, where *Ca*_baseline_ is the basal calcium determined above. The number of IP3R open per peak was defined as the maximum number of IP3R open simultaneously during peak duration. Puffs were defined as calcium events resulting from the cooperation of more than one IP3R. In our spatially-explicit simulations, a calcium signal was considered to be a puff if more than one IP3R were open during the peak and if the average distance traveled by calcium within the duration of this peak was larger than the distance between the simultaneously open IP3R molecules.

### Experimental methods

All experiments were performed as described in [80]. We give below the main outlines of the methods. All experimental procedures were in accordance with the European Union and CNRS UMR5297 institutional guidelines for the care and use of laboratory animals (Council directive 2010/63/EU).

#### Organotypic hippocampal slice cultures

Organotypic hippocampal slices (Gähwiler type) were dissected from 5-7-d-old wild-type mice and cultured 5-8 week in a roller drum at 35°C, as previously described [81].

#### Viral infection

AAV9-GFAP-GCaMP6s [82] was injected by brief pressure pulses (40ms; 15 psi) into the stratum radiatum of 2-3-week old slices from Thy1-YFP-H (JAX:003782) mice 4-6 weeks prior to the experiment.

#### Image acquisition

For Ca^2+^ imaging, we used a custom-built setup based on an inverted microscope body (Leica DMI6000), as previously described in [83]. We used a 1.3 NA glycerol immersion objective equipped with a correction collar to reduce spherical aberrations and thereby allow imaging deeper inside brain tissue [84]. The excitation light was provided by a pulsed diode laser (l = 485 nm, PicoQuant, Berlin, Germany). The fluorescence signal was confocally detected by an avalanche photodiode (APD; SPCM-AQRH-14-FC; PerkinElmer). The spatial resolution of the setup was around 175 nm (in x-y) and 450 nm (z). Confocal time-lapse imaging (12.5 x 25 *μ*m, pixel size 100 nm) was performed at 2Hz for 2.5 min in artificial cerebrospinal fluid containing 125 mM NaCl, 2.5 mM KCl, 1.3 mM MgCl_2_, 2 mM CaCl_2_, 26 mM NaHCO_3_, 1.25 mM NaH_2_PO_4_, 20 mM D-glucose, 1 mM Trolox; 300 mOsm; pH 7.4. Perfusion rate was 2 mL/min and the temperature 32 °C.

#### Image analysis

Spontaneous calcium events were detected and analyzed automatically by ImageJ plugin LC_Pro [85] and then manually verified using Igor Pro (Wavemetrics) [35].

### Statistical analysis

For stochastic models, we generated 20 simulations (with different random numbers) and quantified these simulations as mean ± standard deviation over those 20 simulations. One-way ANOVA was performed to investigate the effect of a given parameter on calcium dynamics. Comparison between two simulation conditions were performed with unpaired Student T test if values followed a Gaussian distribution. Otherwise, a Mann-Whitney test was performed. The same method was applied to compare simulation to experimental results. * is for *p* ≤ 0.05, ** for *p* ≤ 0.01, *** for *p* ≤ 0.001.

## Results

### Spontaneous oscillations in the model

We first analyzed our particle-based model for the CICR signaling system of figure 1. To that end, we compared Monte-Carlo simulations of the particle-based model in two dimensions with the corresponding Mean-Field and Gillespie’s SSA models (see Methods section). Those three models represent three different levels of approximation: the Mean-Field model assumes deterministic kinetics and perfect mixing; the SSA model keeps the perfect mixing hypothesis but assumes stochastic kinetics while the particle-based model assumes stochastic kinetics but accounts for potential non-perfect mixing, i.e. diffusion effects. For comparison with SSA, we first considered perfect mixing of Ca^2+^ ions and IP3 molecules in the particle-based model by setting the diffusion coefficients *D*_Ca_ = *D*_IP3_ = ∞ (see Method section).

Figure 2A shows one simulation sample for each model. A first result is that the stochastic models (SSA and particle-based) do exhibit spontaneous calcium peaks with the parameters of this figure. On top of a background level of roughly 50 Ca^2+^ ions, with fluctuations of roughly ± 20 ions, large and fast peaks arise spontaneously with a total amplitude between 20 and 120 ions above the baseline. In strong opposition, the (deterministic) mean-field model does not show these oscillations: one gets a stationary trace, that systematically coincides with the baseline level of the stochastic traces (Fig 2B). Comparing the two stochastic models (SSA and particle-based) indicates that both display the same basal calcium level (Fig 2B) and the same frequency and mean peak amplitude (Fig 2C). A Piecewise-Deterministic Markov Process (PDMP) model of the system of Figure 1, in which Ca and IP3 dynamics obeyed to the mean-field model while IP3R state dynamics was expressed with SSA, also exhibited robust spontaneous Ca^2+^ peaks (not shown). Altogether, this suggests that stochasticity is necessary for spontaneous calcium signals to occur in this model.

**Fig 2.**
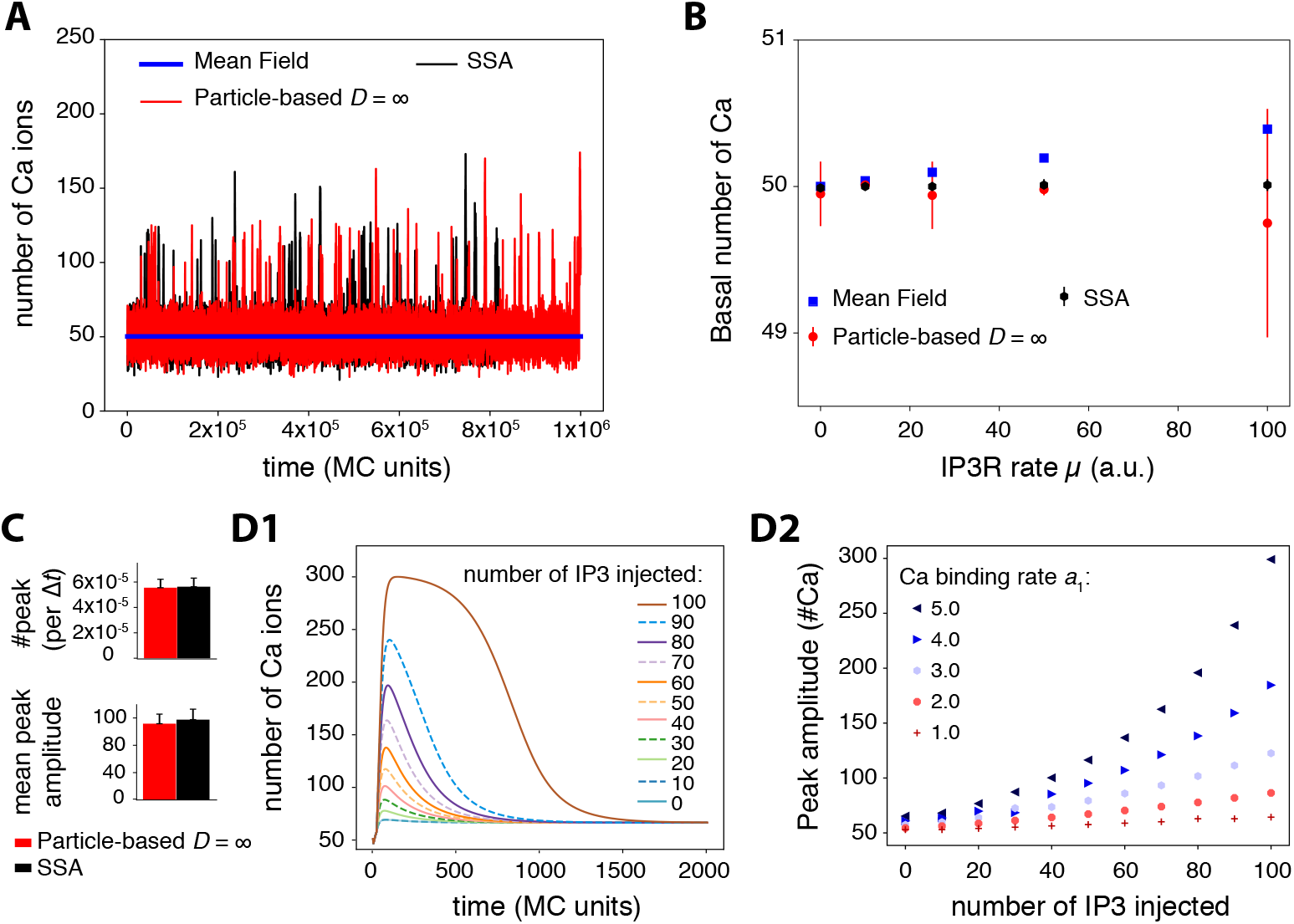
Model exploration. (*A*) Spontaneous transients are observed in simulations of the particle-based and the Gillespie’s SSA model but not in the Mean Field model. (*B*) The three models display the same basal calcium level when *μ*, the calcium influx rate through open IP3R channels, increases. The higher variability in the stochastic models reflects the integer value of basal calcium (either 49 or 50, depending on simulations). (*C*) Quantification of calcium transients in the stochastic models (calcium peak frequency and mean peak amplitude). No significant difference between the two models was observed. (*D*) Excitability of the Mean-Field model: increasing quantities of exogenous IP3 molecules were injected at time *t* = 20Δ*t*, after model equilibration. The amplitude of the resulting calcium response (*D1*) was quantified depending on the amount of IP3 injected and the value of the binding rate constant to the first calcium IP3R site, *a*_1_ (*D2*). Parameter values for the particle-based model: *D*_Ca_ = *D*_IP3_ = ∞ (perfect mixing) and *η* =1, *R_γ_* = 200, i.e. no IP3R channels clustering, and no co-localization of IP3R with IP3R-independent Ca^2+^ sources. For SSA and particle-based models, the figure shows the average ± standard deviation over 20 simulations.

We next searched for the dynamical mechanism that gives rise to those spontaneous peaks. A thorough numerical parameter exploration of the mean-field model failed to evidence the existence of Hopf bifurcations or of any other bifurcation that would generate limit-cycle oscillations in the model (not shown). This is a distinctive feature of our model, since spontaneous oscillations in the vast majority of IP3R-mediated calcium signaling models arise from limit-cycle generating bifurcations [48–50]. This is however not unexpected since the simplifications made to derive our model significantly reduced its nonlinearity compared to these models, and the emergence of limit-cycle bifurcations demands strong nonlinearity. For instance, limit-cycle oscillations in the classical Li and Rinzel model [50] disappear when IP3R opening needs less than three open monomers (not shown). However, our model retains enough nonlinearity to exhibit excitability. To evidence this, we used the mean-field model, waited until all concentrations reached their stationary state, and injected an increasing amount of exogenous IP3 molecules. In response to this IP3 injection, a calcium transient was obtained, before relaxation to the stationary state (Fig 2D). Figure 2 D2 shows how the resulting transient amplitude depends on the amount of injected IP3. For low values of IP3R calcium binding rate (first site), *a*_1_, the calcium response is basically linear with the number of injected IP3: doubling the amount of IP3 injected only doubles the amplitude of the calcium response. However, as a1 increases, peak amplitude becomes a strongly nonlinear function of the number of IP3 injected. With *a*_1_ = 5 a.u. for instance, doubling the number of injected IP3 from 50 to 100, results in an almost threefold increase of the calcium response. Therefore the mean-field model with large values of a1 is an excitable system that amplifies the fluctuations of IP3 in its calcium responses. We conclude that spontaneous calcium transients occur in the system of figure 1 through the interplay of the stochasticity of the SSA or particle-based models and the underlying excitability of the system.

### Transitions between calcium activity regimes

The experimental and modeling literature on intracellular calcium signals distinguishes two classes of localized calcium peaks: puffs and blips [86]. Blips refer to brief and weak peaks that correspond to the opening of a single IP3R channel (or a single IP3R channel tetramer), whereas puffs are longer and higher peaks resulting from the concerted opening of a group of nearby IP3R channels (or tetramers thereof), via the calcium-induced calcium-release principle. We next examined whether our model was able to reproduce these observations.

We carried out parameter exploration of the particle-based model in conditions of perfect mixing for mobile molecules (*Ca* and *IP*3) and uniform spatial distribution of the immobile ones (PLC*δ*, IP3R). As expected, we found that calcium peaks frequency depends on parameter values (Fig 3A). When the rate of calcium influx through open IP3R channels *μ* or the binding rate constant to the first Ca IP3R site *a*_1_ are too small, the model does not exhibit calcium peaks at all, only fluctuations around a stationary state (Fig 3C★). This is in agreement with our analysis of the systems excitability above, that evidenced excitability only for large enough values of *a*_1_ (Figure 2D2). Note however that in the model, IP3R openings do not necessarily lead to a calcium peak, especially for low values of both *μ* and *a*_1_ (Fig 3C★). Spontaneous calcium transients are obtained in the particle-based model beyond threshold of (*μ, α*_1_) values, with a peak frequency that increases with parameters values (Fig 3A). Inspection of the maximal number of open IP3R per peak reveals that not only the frequency, but also the type of these transient signals changes with parameters values (Fig 3A): the less frequent signals are generally associated with a single open IP3R per peak (Fig 3C▬), corresponding to blips, whereas the high-frequency spontaneous signals rely on the opening of 2 − 12 IP3R in a peak (Fig 3C•), corresponding to puffs. In agreement with experimental observations [68,87], calcium puffs in the particle-based model are characterized by higher peak amplitude and peak duration compared to blips. Taken together, these results show that our particle-based model not only reproduces the existence of spontaneous calcium peaks in conditions of low copy numbers, it is also able to reproduce the existence of different types of localized calcium transients, in agreement with experimental measurements.

**Fig 3.**
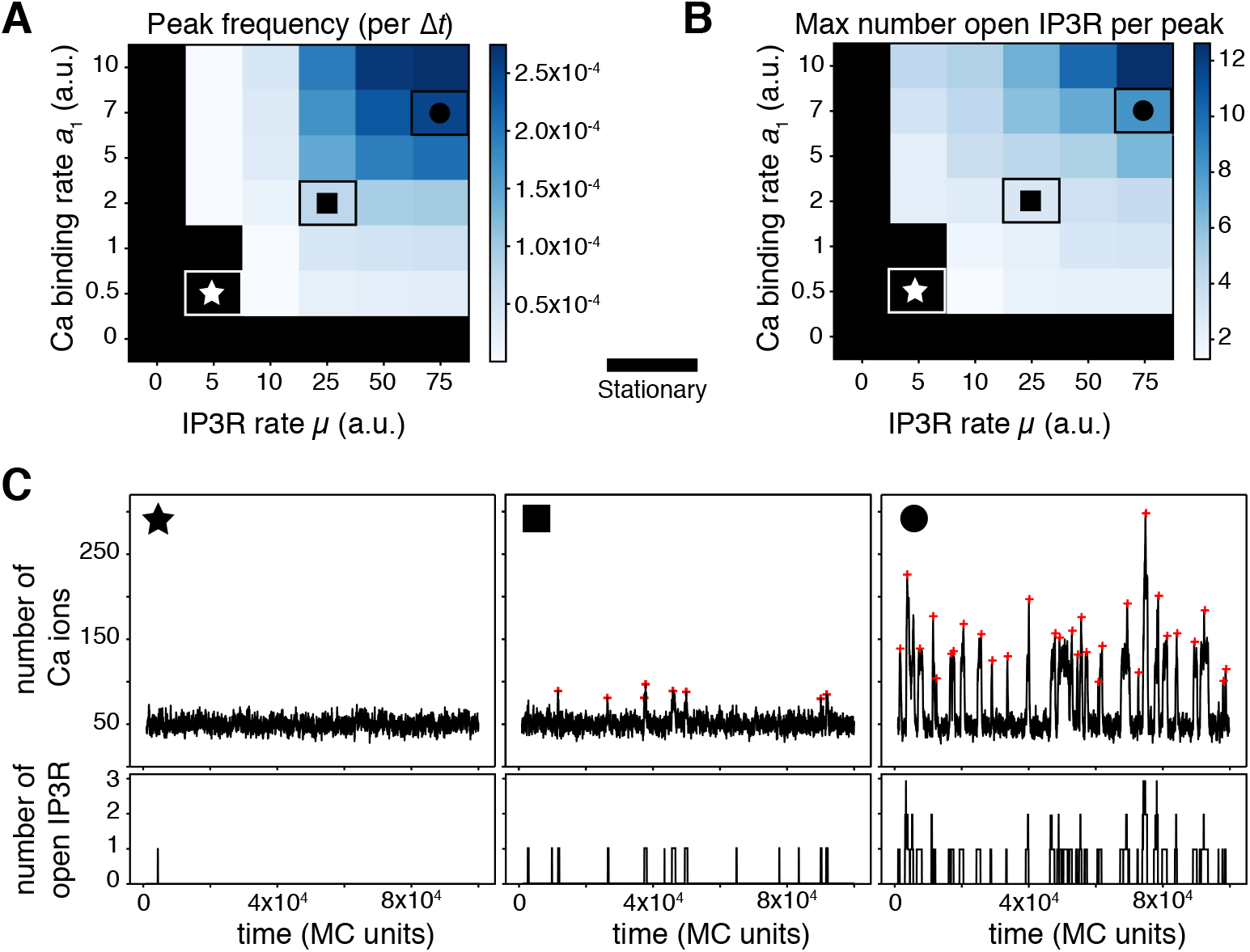
The particle-based model produces different calcium activity regimes depending on parameter values. Color-coded map of variation of the peak frequency, expressed as the number of calcium peaks per MC time step (*A*) and as the maximal number of IP3R channel open per peak (*B*). The color scale is given for each map. The black area corresponds to the stationary regime. Note that the x and y-axis scales in (A) and (B) are not regularly spaced. The symbols ★, ∎ and • locate parameter pairs that are illustrative of the three dynamical regimes shown in (C): stationary (★, *μ* = 5, *α*_1_ = 0.5), blips (∎, *μ* = 25, *a*_1_ = 2) and puffs (•, *μ* = 75, *α*_1_ =7). Red crosses show the locations of peaks from automatic detection. *D*_Ca_ = *D*_IP3_ = ∞, *η* = 1, *R*_γ_ = 200.

### Impact of calcium diffusion coefficient on calcium signals

A modeling study has demonstrated the necessity to account for the stochasticity inherent to calcium diffusion when modeling calcium signaling in small volumes [88]. We next investigated the impact of calcium diffusion on calcium dynamics in the particle-based model. In neurons or astrocytes, the amount of endogenous calcium buffers is large so that the diffusion distance of free calcium is believed to be very small. Many of the endogenous buffers are however mobile. Therefore, our diffusion coefficient for calcium is to be interpreted as an effective diffusion coefficient lumping together calcium buffering by mobile endogenous buffer and diffusion of these buffers.

Moreover, several plasma membrane proteins, in particular the Na^+^-Ca^2+^ exchanger (NCX) have been observed to co-localize with ER proteins in neurons and astrocytes [89]. Such a co-localization of calcium signaling molecules might imply spatial organizations including raft-like micro-domains. This organization seems essential for calcium wave propagation in astrocytes [90]. Moreover, mGluR5-ER proteins co-clusters mediated by an interaction with Homer1 scaffold protein have been observed in astrocytic processes [91]. Homer1 is also known for increasing calcium activity in neurons by increasing IP3R-mGluR5 proximity [92]. Those experimental studies suggest that several calcium sources are co-localized with ER proteins in astrocytes and that it might alter calcium dynamics. Such a co-localization could be crucial for calcium signaling, in particular in small volumes. We thus placed our study of the influence of calcium mobility on calcium signaling in a framework where calcium sources (IP3R-dependent and IP3R-independent) can co-localize.

To this end, the IP3R-independent calcium influx in the cytosol (from e.g. plasma membrane transporters or channels) was made dependent on parameter *R*_γ_, that sets the distance from IP3R receptors within which new calcium ions are injected in the cytosol when they originate from IP3R-independent fluxes (see Methods section). When *R_γ_*=0, the initial location of the new calcium ion is shared with an IP3R channel whereas when *R_γ_* increases, the injection positions of new calcium ions are increasingly uncorrelated from those of the IP3R channels. When *R_γ_* becomes as large as the size of the reaction surface (i.e. for *R_γ_* =100), the injection position of the new calcium ion is effectively independent of the positions of the IP3R channels.

Our simulations show that the impact of the calcium diffusion coefficient is mainly visible when calcium sources are co-localized, i.e. for small values of *R_γ_*. Figures 4A-B compare a representative simulation obtained when Ca^2+^ diffuses slowly (A) with a simulation obtained with perfectly-mixed calcium (B), in a case where the IP3R receptors are not clustered (*η* = 1). Those representative simulations hint that the peak frequency is much larger with slow calcium, and suggests that slow calcium diffusion slightly favors the puff regime compared to perfect mixing. The systematic quantifications of figure 4C-D confirms these interpretations: when IP3R-dependent and IP3R-independent calcium sources are co-localized, i.e. for *R_γ_* < 5, the value of *D*_Ca_ controls calcium transient frequency, as well as the probability to observe a puff. The effects are strong: for instance for *R*_γ_ = 0, decreasing *D*_Ca_ from 5 to 0.1 increases the frequency roughly threefold. However, when the IP3R-independent influx was not co-localized with IP3R channels (i.e. for *R_γ_*≥ 5), both the peak frequency and the type of signal were found not to depend on the calcium diffusion coefficient anymore. Those results suggest that calcium diffusivity could control the frequency and type of calcium signals within astrocytes when IP3R channels are co-localized with IP3R-independent calcium sources.

**Fig 4.**
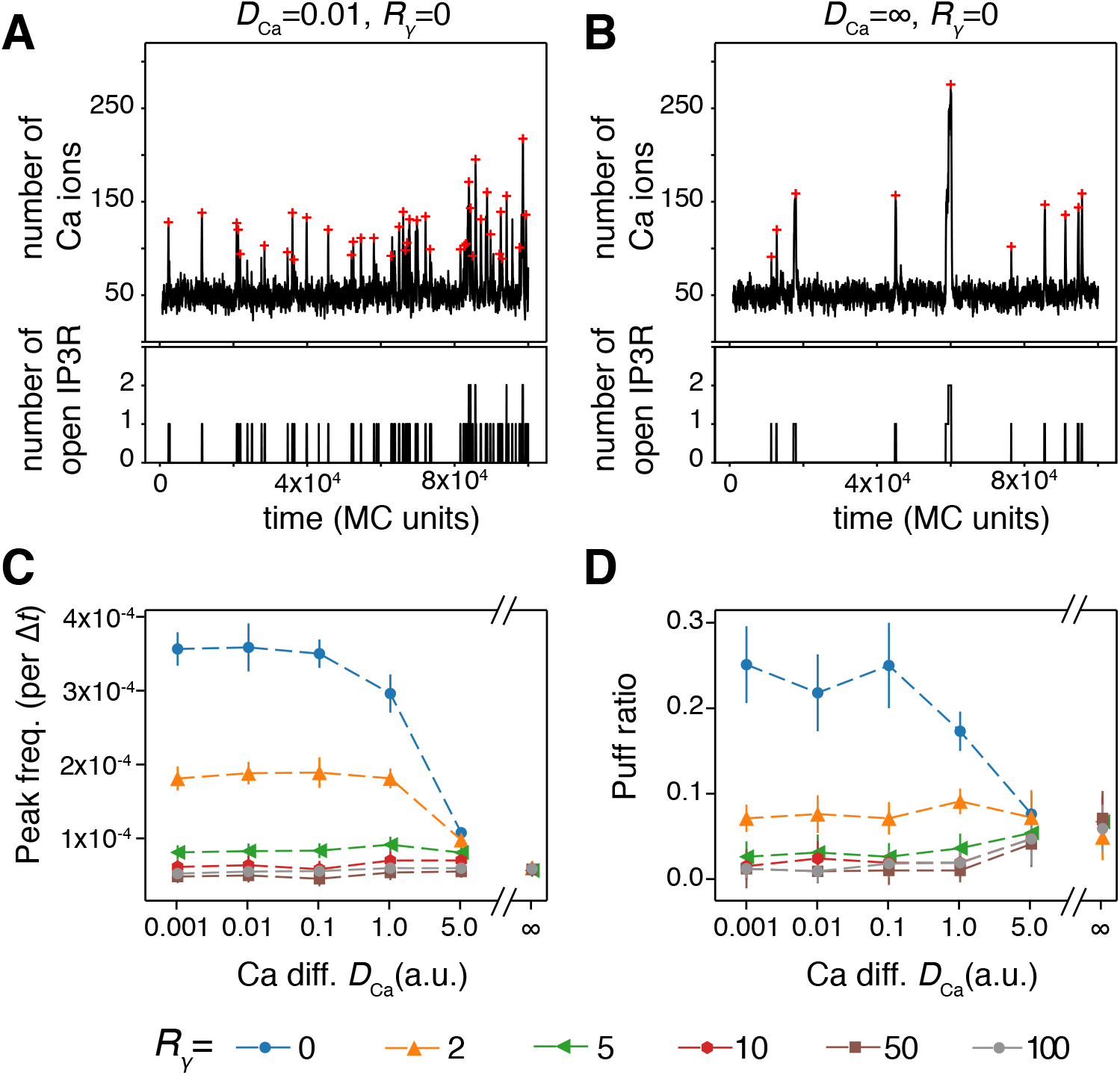
Ca^2+^ diffusion modulates the temporal characteristics of the signals upon co-localization. Representative simulations of the particle-based model showing both calcium trace and number of open IP3R for co-localized calcium sources (*R_γ_* = 0) in the case of slow calcium diffusion (*A*) or perfect-mixing of calcium (*B*). The red crosses show peak locations from automatic detection. The impact of calcium diffusion coefficient *D*_Ca_ on peak frequency (*C*) and the amount of puff (*D*) are shown for different values of the co-localization parameter *R_γ_*: from *R_γ_* = 0 (IP3R are not clustered but co-localized with other calcium sources) to *R_γ_* = 100 (IP3R are neither clustered nor co-localized). The puff ratio quantifies the fraction of peaks that are puffs. Data are presented as mean ± standard deviation over 20 simulations. Lines are guide for the eyes. Note that the x-axis scale in (*C*) and (*D*) are not regularly spaced. Other parameters: *η* = 1 (no clustering), *a*_1_ = 1, *μ* = 50.

### IP3R clustering controls calcium signals when co-localized

Experimental data demonstrate that IP3R in SH-SY5Y and COS7 cells are not uniformly distributed on the ER membrane but form clusters [68,87]. We next investigated the impact of IP3R clustering on calcium signal dynamics in our particle-based model. Simulations were performed with *D_ca_*=0.1 and various amounts of co-localization between IP3R channels and other calcium sources (parameter *R_γ_*). Representative simulations for uniformly-distributed IP3R channels (*η* =1) and strongly clustered IP3R (*η* = 50) are presented in figure 5A-B. In these two examples, the IP3R were weakly co-localized with the IP3-independent calcium sources (i.e. *R_γ_* = 10). These traces indicate that the frequency and type of calcium signal in this case is heavily dependent on the spatial distribution of IP3R channels: clustered IP3R seem to exhibit much larger peak frequency and slightly more frequent puffs. However, here again this effect is quite mitigated by the amount of co-localization between IP3R channels and the IP3R-independent calcium sources. In particular, the dynamical range of the modulation by IP3R cluster size *η* (i.e. the ratio between the frequency at *η* = 50 and *η* = 1) is maximal for intermediate co-localizations (2 ≤*R_γ_*≤ 10) but the calcium peak frequency is hardly dependent on η when co-localization is either very strong (*R_γ_*< 2) or very weak (*R_γ_*≥ 50). Increasing clustering also tends to improve the emergence of puffs, although the effect is significant only for strong co-localization (*R_γ_*≤ 2, Fig 5D). We emphasize that in such cases of strong co-localization, the regime of calcium activity (puffs *vs* blips) changes by simply rearranging the spatial distribution of the IP3Rs, without changing any of the kinetics parameters of the model.

**Fig 5.**
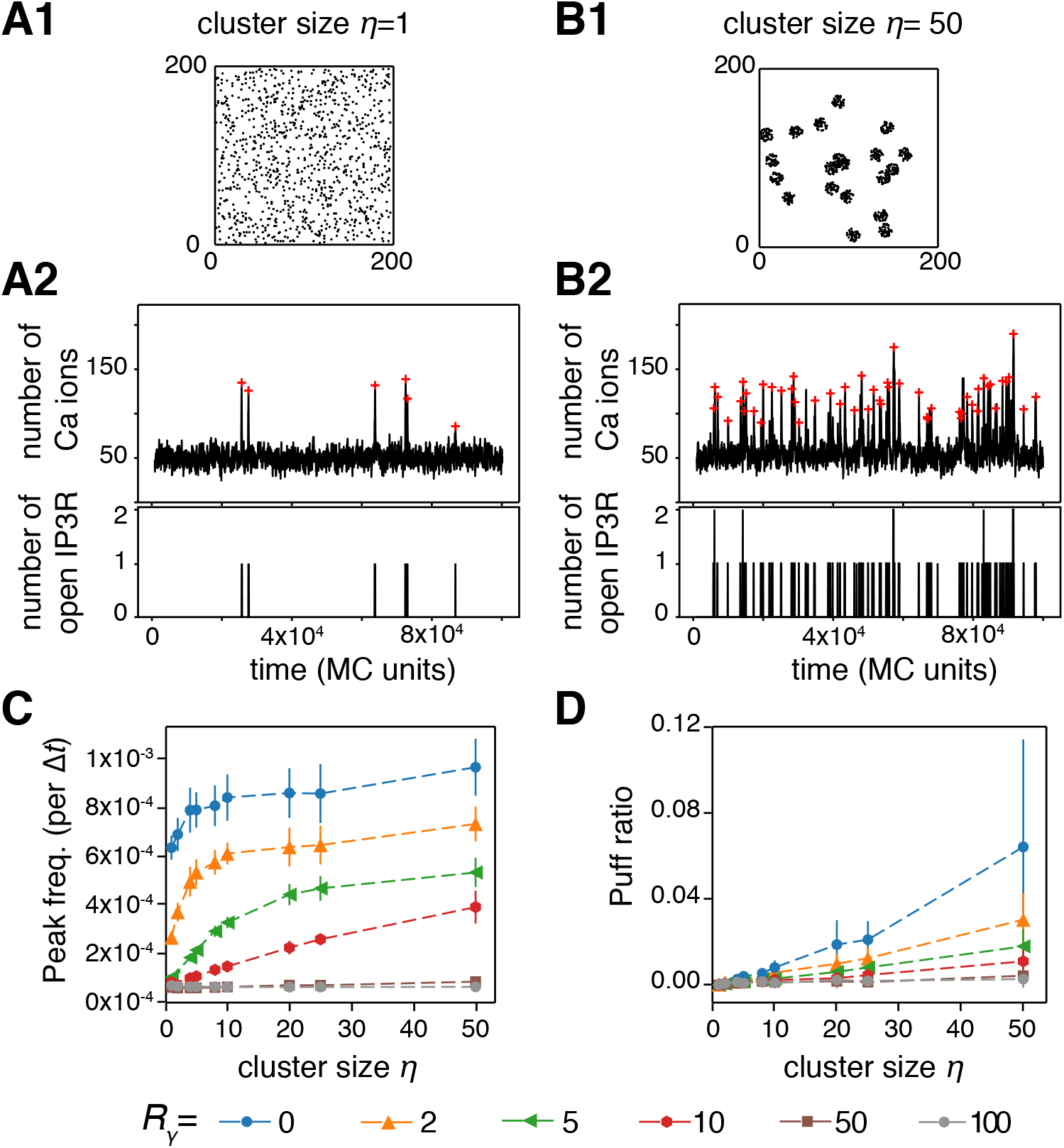
IP3R clustering modulates calcium signals when co-localized. Representative simulations of the particle-based model with the corresponding IP3R distribution over space, the calcium trace and number of open IP3R for weakly co-localized calcium sources (*R_γ_* = 10) in the case of uniform distribution of the IP3R (*A*) or strongly clustered IP3R (*B*) are illustrated. The red crosses show peak locations from automatic detection. The impact of IP3R cluster size *η* on calcium peak frequency (*C*) and on the amount of puffs (*D*) are shown for different values of the cluster size: from *η* =1 (IP3R are not clustered) to *η* = 50 (strong clustering). Data are presented as mean ± standard deviation over 20 simulations. Lines are guide for the eyes. Other parameters: *D*_Ca_ = 0.1, *a*_1_ = 1, *μ* = 50.

Taken together, these simulation results pinpoint the interplay between calcium source co-localization and the degree of IP3R clustering as a crucial modulator of temporal characteristics of the calcium signals and of the signaling regime. In particular, they suggest that in the presence of certain amount of co-localization between IP3R channels and other sources of calcium influx in the cytosol the spontaneous calcium peak frequency can have a large amplitude variation. Within this range of parameters, calcium peak frequency can be finely tuned by the geometry of the colocalization.

### Simulations in a compartmentalized 3d geometry reproduce spontaneous calcium microdomains signals

The above 2d simulations of the particle-based model have the advantage of a good computational efficiency, which makes them suitable for parametric studies with averaging over a number of Monte-Carlo simulations. However, the 2d setting does not facilitate the comparison of the copy number of molecules in the simulations with species concentrations as measured experimentally. Moreover, it is difficult to investigate with a 2d setting the impact of the fact that IP3R channels are specifically localized at the surface of the ER membrane and not freely diffusing in the cytosol bulk. To tackle those questions, we carried out simulations of our model in a more refined three-dimensional setting (Fig 6), in which we could adjust more precisely molecule concentrations, reaction volume and cytosol compartmentalization to what is expected in fine astrocytic processes. We then compared our simulations to experimental measurements of calcium dynamics in microdomains of comparable dimensions in mice hippocampal organotypic culture (Fig 6A).

**Fig 6.**
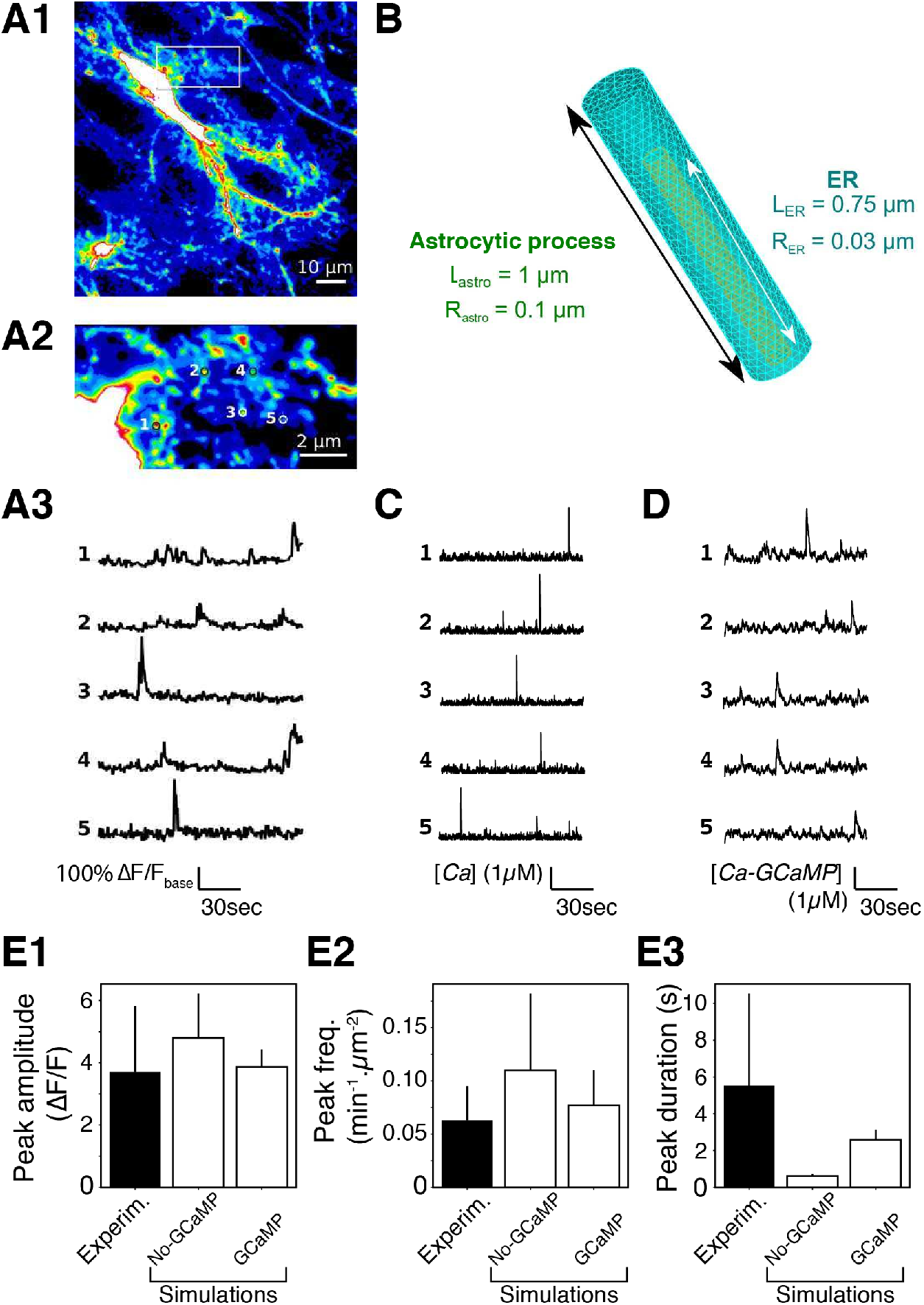
3d model simulations in fine astrocyte processes successfully reproduce calcium microdomains signals. (*A*) Experimental monitoring of the spontaneous local Ca^2+^ signals in astrocytic sponge-like processes. Panel *A1* shows a ‘summed projection’ of a confocal time lapse image stack of a GCaMP6s-expressing astrocyte. Panel *A2* illustrates magnification of the boxed region of panel *A1*. Panel *A3* displays spontaneous calcium traces from the regions of interest shown in (*A2*). (*B*) The 3d geometry used for the 3D model is a cylinder of length L_astro_=1 *μm* and radius R_astro_=0.1 *μm*, with ER as a thinner cylinder inside. The interior volume is roughly 0.03 fL. (*C*) Representative simulations of calcium dynamics within the above cylinder with the “No-GCaMP” simulations. The raw signal corresponds to cytosolic free calcium concentration. (*D*) Representative simulations of calcium dynamics within the above cylinder with the “GCaMP” simulations. The raw signal corresponds to calcium-bound GCaMP concentration. For both simulation types, parameter values were partly taken from the literature and partly adjusted for fitting calcium traces shown in *A* (reported in Table 2). (*E*) Quantitative comparisons of the spontaneous calcium signals measured experimentally (black bars) or simulated with the “No-GCaMP” or “GCaMP” models (white bars). The compared quantities are peaks amplitude in terms of Δ*F/F* ratio (E1), their frequency (measured in *min*^−1^ for each *μm*^2^ area, E2) and duration E3.

As 80% of calcium activity occurs in astrocytic ramifications that cannot be resolved by optical microscopy [40], astrocytic calcium signaling models must take into account small volumes associated to it. For that purpose, we created the 3d structure mimicking one process geometry shown in Figure 6B. The reaction volume was chosen to match the range of sizes that are within reach of current imaging methods: a 1 *μm*-long cylinder of 100 nm radius (i.e. a volume around 0.03 fL), inside which we position a 0.75 *μm*-long cylindrical ER with a radius of 30 nm. In this 3d implementation, calcium and IP3 molecules diffuse in the bulk 3D space located between the external (plasma) membrane and that of the ER, while IP3R molecules are distributed uniformly at random over ER membrane surface.

Our calcium imaging of the calcium dynamics in fine astrocyte processes reveals the sponge-like structure of the processes 6A1, with localized submicron calcium microdomains (regions of interest (ROI) in figure 6A2) of size that can be less than 0.5*μm*^2^. The corresponding calcium traces display infrequent (a few hundredths of Hz) peaks with average amplitude around 4 (Δ*F/F*) and typical duration of ≈ 5 seconds (fig 6A3 and E). Notice that these experimental traces correspond to spontaneous signals to the extent that they were measured in the absence of any neuronal nor astrocytic stimulation. In particular, TTX application in this preparation did not alter peak frequency [80].

Our first noticeable result is that our model is able to reproduce the emergence of spontaneous calcium peaks of comparable frequency and signal-to-noise ratio (fig 6C).

This result therefore indicates that spontaneous calcium signals can emerge in the fine processes even with a realistic basal calcium concentration of 83 ± 29 nM, which corresponds to only one to two (1-2) calcium ions in the whole cylinder. However, quantification of the signal properties (fig 6E) shows that the agreement is only qualitative, since the peak amplitude and frequency are slightly too large while peak duration is much shorter than in the experiments (fig 6E, “No-GCaMP” simulations). Adding GCaMP6s to the model improved drastically the match between simulations and experimental data (fig 6D). The properties of the signals now do match experimental data (fig 6E, “GCaMP” simulations). In particular, peak duration is much larger than without GCaMP (fig 6E3). Note that our experimental statistics are tightly associated with the temporal sampling frequency used in the experiments (2 Hz) since very fast calcium events may be accessible only to higher sampling frequencies [40]. In particular, the experimental peak frequency measured might have been higher with better temporal resolution. In any case, our results show that genetically encoded calcium indicators (GECIs), such as GCaMP6s, may change local calcium concentration, in particular close to open IP3R channels, leading to an increased peak duration.

Interestingly, the well-mixed implementation of this model in the cylinder geometry, with the same parameter values, displays a higher peak frequency (data not shown). This reflects the importance of stochasticity and local variations of calcium concentration for tuning IP3R activity and thus the overall calcium signal dynamics. Together these results demonstrate that our model, without any endogenous buffers, is enough to reproduce calcium signals within fine astrocytic processes in a quantitative way, making it a powerful tool to investigate calcium dynamics in the small volumes associated with the PAPs.

## Discussion

Recent experimental reports suggest that the complete dependence of cytosolic calcium transients on IP3R2 is only observed in the astrocyte cell body whereas calcium signals measured within astrocytic processes are a mix of IP3R2-dependent and non-IP3R2-dependent calcium signals [21, 30]. The identity, subtype and localization of the receptors responsible for non-IP3R2-dependent calcium signals in astrocytes, in particular their processes, are still to be uncovered. However, our study sheds light on the importance of the localization of these various calcium sources. Our simulation results indeed indicate that whenever IP3R channels are (even moderately) co-localized with IP3R-independent calcium sources, e.g. plasma membrane calcium channels, the degree of IP3R clustering and/or the mobility of the calcium buffers will have a strong impact on the frequency and amplitude of the spontaneous calcium signals. In particular, our simulations predict that two astrocyte processes expressing exactly the same repertoire of channels, pumps and receptors but in a different spatial organization (for instance various degrees of clustering or co-localization), can exhibit very different types and properties of spontaneous calcium events. This could result in significant variability of the calcium response of different processes, even from the same cell.

During the past few years, fine astrocytic processes have been regarded as devoid of ER [93,94]. This questions the validity of our model, in which the presence of ER-attached IP3R in the process is crucial for spontaneous activity. We however note that a recent EM study has observed that ER dynamically ramified in astrocyte perivascular processes *in vivo* and detected contact sites between ER processes and plasma membrane, often positioned in apposition to neuronal synapses [95]. Such contiguous membranous juxtapositions would definitely validate the presence of ER in PAPs. Although dynamical ER remodeling has been reported in dissociated astrocyte culture [96], technical limitations have prevented direct investigation of ER localization within PAPs *in vivo* or in slices. To our knowledge, it is not even clear whether astrocytic ER is continuous or consists in several independent reservoirs. Super-resolution microscopy of cellular ER and mitochondrial dynamics and structure (resolution ≈ 100nm) has recently been developed and could help solve the controversy regarding the presence of ER in fine processes [97,98]. Correlative super-resolution fluorescence imaging and electron microscopy approaches can yield a resolution of less than 50 nm (down to 10nm) [99], which is very promising avenue to PAPs ultrastructure investigation. At any rate, since the IP3R pathway is involved in calcium dynamics, further investigations regarding ER sub-cellular localization, sub-compartmentalization and dynamics are crucial for better understanding astrocyte information processing. Meanwhile, a straightforward extension of our computational model would be to simulate neuronal stimulation-triggered calcium dynamics.

IP3Rs are thought to assemble as tetramers, and a recent experimental study suggested that the four subunits of the tetramer must be simultaneously bound to IP3 for the tetramer to allow calcium influx, independently of cytosolic calcium or ATP concentrations [100]. Actually, the original IP3R models predicted that subunit cooperativity for calcium binding is also necessary to fit experimental data of IP3R dynamics [48,50]. Even though the IP3R binding sites for calcium have been characterized, their roles in IP3R dynamics are still poorly understood [101]. The requirement for inter-subunit cooperativity, in which the 4 IP3 binding sites should simultaneously be bound for the tetramer to open, is expected to hinder the emergence of spontaneous calcium events. In a subset of simulations, we have replaced our non-cooperative IP3R model, in which the binding of a single IP3 site is enough to open the monomer channel, with the cooperative model proposed by Bicknell and collaborators [102]. With 100 nM basal IP3 and Ca^2+^ [65,67], we could not produce spontaneous calcium signals in these conditions, even after a search of the parameter space to locate parameters allowing spontaneous activity with this cooperative model. This result casts doubts on the reality of spontaneous calcium signals in astrocytes when the basal IP3 and Ca^2+^ concentrations are of the order of 100 nM. A number of studies have reported higher calcium concentration localized at the vicinity of calcium channels [71,103,104]. Such calcium microdomains, in the vicinity of IP3R, could facilitate the emergence of spontaneous signals from cooperative IP3Rs in thin processes.

On the other hand, experimental evidence for spontaneous calcium signals in astrocytes is still debated. Even in the absence of presynaptic neural activity, presynaptic axon terminals do probabilistically release neurotransmitter vesicles, generating so-called miniature EPSCs. Bafilomycin application has been used in several experimental studies to investigate the dependence of astrocytic calcium signals on EPSCs, because this inhibitor of V-ATPases inhibits miniature EPSCs by blocking the refill of presynaptic vesicles. However, the impact of bafilomycin bath application on the frequency of spontaneous calcium signals in astrocytes has proven variable (compare e.g. [105] and [106]). In our preparation, bath-application of bafilomycin strongly decreased peak frequency and amplitude [80]. As bafilomycin has a wide range of effects on calcium signaling that is independent of its effect on the refill of presynaptic neurotransmitter vesicles [107,108], we cannot conclude whether those signals are triggered by EPSCs and further investigation is needed to decipher whether the ‘‘spontaneous” calcium signals reported in astrocyte processes are due to spontaneous release of presynaptic vesicles or rely on a synapse-independent mechanism inherent to the CICR system.

For simplicity, IP3R clustering in our model was considered static during simulation time. Experimentally, though, IP3R clustering might be highly dynamic [109,110]. Several molecules can trigger IP3R clustering, including IP3 and calcium themselves [109,110], through a mechanism that may include the lateral diffusion of IP3R on the ER surface [110] or be independent from it [72]. Beyond this IP3R classification into clustered and un-clustered populations, another approach is to quantify single IP3R channels based on their mobility. A recent study on HeLa cells [43] indicates that calcium signals emerge most of the time from immobile IP3R, which are found in apposition to ER-plasma membrane junctions, whereas the mobile IP3R fraction would not be involved in calcium influx. Our simulation results, in agreement with previous IP3R-mediated calcium models [111,112], indicate that IP3R clustering can lead to an increase of the frequency and amplitude of their calcium signals. This result is in contradiction with a previous modeling study that concluded in favor of a reduction of IP3R channel activity upon IP3R clustering [113]. This discrepancy might rely on the different modeling choices. In particular, the model in [113] incorporates a 5-state IP3R model derived from ref [114,115]. All of those modeling studies however agree that dynamical IP3R clustering could be a mechanism used by astrocyte processes to modulate their calcium signals. This could provide astrocyte processes with a capacity for information processing plasticity.

In our model, the value of the rate constant for calcium binding on IP3R changes the type of spontaneous dynamics (e.g. blips vs puffs) in addition to its characteristics (frequency, amplitude). Experimentally, several post-transcriptional mechanisms can modulate IP3R affinity. For instance, phosphorylation of type-1 and −2 IP3R by cAMP-activated PKA increases the affinity of IP3R to calcium and IP3 [116]. At a larger time scale, the sensitivity of IP3R to calcium is encoded in a sequence of calcium sensor (Cas) region that differs depending on the IP3R isoform [114,117,118]. Since multiple IP3R isoforms seem to be involved in calcium signaling within astrocytic processes [35], they could assemble into a variety of homo-or hetero-IP3R tetramers that would exhibit a range of calcium and IP3 affinity. In addition, immobile or weakly mobile endogenous calcium buffers are responsible for an effective intracellular calcium diffusion that is an order of magnitude slower than free calcium ions [74]. The variability of these endogenous calcium buffers, with various kinetics and various diffusion coefficients, is large [119]. Some of them are overexpressed in hippocampal and striatal astrocytes, possibly in a region-specific pattern [120]. Our simulation results indicate that the value of the effective Ca^2+^ mobility also participates in the determination of the type and characteristics of the spontaneous events, thus confirming previous modeling approaches [88]. Such a regional differential expression of the genes coding for endogenous calcium buffers could therefore be involved in the regional variability of astrocytic calcium signalling [121]. Therefore, the IP3R repertoire, the post-transcriptional regulation of IP3R affinity and the differential expression of endogenous calcium buffers could also be potential determinants allowing a range of responsiveness and spatio-temporal characteristics of calcium signals in astrocyte processes.

To conclude, we have presented a spatially-explicit stochastic model to investigate intracellular calcium signaling based on CICR in small sub-cellular volumes. Two recent studies proposed models for the simulation of astrocytic calcium signals in 3d with deterministic differential equation models that correspond to cellular volumes large enough to validate a law of large numbers [94, 122]. To our knowledge, our model is the first model suited to reproduce spontaneous calcium signals in the finest astrocyte processes, where low copy number and spatial localization effects are expected to be more prominent than in larger volumes. Since these fine processes are thought to be the place of initiation of neuron-astrocyte interactions, we believe this model might be useful to investigate the initiation and spatiotemporal integration of calcium signals in the sponge-like network of the astrocyte process, a prerequisite to understand neuron-astrocyte communication.

## Acknowledgments

We thank Iain Hepburn, Weiliang Chen and Andrew Gallimore of the Computational Neuroscience Unit, OIST, Okinawa, Japan for discussion about 3d meshes and STEPS.

