## Supplemental Table S1 for "Simulation of calcium signaling in fine astrocytic processes: effect of spatial properties on spontaneous activity"

**Table S1. Parameter values and initial conditions for the 3d model without GCaMP**

| Parameter | Description | Value | Ref. |
|---|---|---|---|
| V | Cell Volume | $2.81 \cdot 10^{-17}$ L | [1] |
| | IP3 dynamics | | |
| $IP_0$ | Initial IP3 number/conc | 3 molec. i.e 177 nM | [2] |
| $D_{\mathrm{IP3}}$ | IP3 diffusion | 223 $\mu$m$^2$.s$^{-1}$ | [3, 4] |
| $N_{\mathrm{plc}}$ | PLC$\delta$ number/conc. | 1696 molec. i.e 100 $\mu$M | [5] |
| $\delta$ | PLC$\delta$ max rate | 1 s$^{-1}$ | - |
| $\beta$ | IP3 decay | $1.2 \times 10^{-4}$ s$^{-1}$ | - |
| | $Ca^{2+}$ dynamics | | |
| $Ca_0$ | Initial $Ca^{2+}$ number/conc. | 5 molec. i.e 295 nM | [6] |
| $D_{\mathrm{Ca}}$ | $Ca^{2+}$ diffusion | 13 $\mu$m$^2$.s$^{-1}$ | [3] |
| $\mu$ | $Ca^{2+}$ flux through open IP3R | $2.5 \times 10^3$ s$^{-1}$ | - |
| $\gamma$ | cytosolic $Ca^{2+}$ influx | $6 \times 10^{-7}$ s$^{-1}$ | - |
| $\alpha$ | $Ca^{2+}$ decay constant | 5 s$^{-1}$ | - |
| | IP3R | | |
| $N_{\mathrm{IP3R}}$ | IP3R number | 50 | [7] |
| $d_{\mathrm{IP3R}}$ | IP3R interact. distance | 1 triangle | - |
| | IP3R binding | | |
| $a_1$ | First Ca | $1.5{\times}10^6$ M$^{-1}$.s$^{-1}$ | - |
| $a_2$ | IP3 | $1.5 \times 10^6$ M$^{-1}$.s$^{-1}$ | - |
| $a_3$ | Second Ca | $1 \times 10^5$ M$^{-1}$.s$^{-1}$ | - |
| | IP3R dissociation | | |
| $b_1$ | First Ca | 70 s$^{-1}$ | - |
| $b_2$ | IP3 | 70 s$^{-1}$ | - |
| $b_3$ | Second Ca | 70s$^{-1}$ | - |
